# A Microfluidic Platform for Functional Testing of Cancer Drugs on Intact Tumor Slices

**DOI:** 10.1101/2020.03.02.973818

**Authors:** A.D Rodriguez, L.F Horowitz, K. Castro, H. Kenerson, N. Bhattacharjee, G. Gandhe, A. Raman, R. J. Monnat, R. Yeung, R.C. Rostomily, A. Folch

## Abstract

Present approaches to assess cancer treatments are often inaccurate, costly, and/or cumbersome. Functional testing platforms that use live tumor cells are a promising tool both for drug development and for identifying the optimal therapy for a given patient, i.e. precision oncology. However, current methods that utilize patient-derived cells from dissociated tissue typically lack the microenvironment of the tumor tissue and/or cannot inform on a timescale rapid enough to guide decisions for patient-specific therapy. We have developed a microfluidic platform that allows for multiplexed drug testing of intact tumor slices cultured on a porous membrane. The device is digitally-manufactured in a biocompatible thermoplastic by laser-cutting and solvent bonding. Here we describe the fabrication process in detail, we characterize the fluidic performance of the device, and demonstrate on-device drug-response testing with tumor slices from xenografts and from a patient colorectal tumor.

## Introduction

The average cost of developing a new cancer drug is now more than $650 million.^1–3^ Unfortunately, most pharmaceutical drugs in clinical development phase never make it to market; in a recent report, 76% of clinical failures were due to a lack of efficacy (52%) or to safety issues (24%), with almost 30% being cancer drugs.^4,5^ One of the main causes of this expensive gridlock is that pre-clinical animal tests do not accurately predict toxic doses and drug metabolism observed in humans.^6^ A recent study of clinical drug development success rates indicated that oncology had the lowest likelihood of approval from phase 1.^4,5^ Clearly, there is an urgent need for better functional drug assays based on human tissue which would more closely mimic patient disease and predict clinical outcomes to complement ongoing efforts using 3D cell culture systems and animal models. Also, functional drug assays on patient tissue have been proposed as a way to guide therapy decisions to complement present genomic approaches to personalized medicine.^7^ Personalized approaches could lower treatment toxicity, improve patient’s quality of life, and, most importantly, reduce mortality.

With increasing evidence that tumor-associated stromal cells play key roles in tumorigenesis and tumor progression,^8^ it has become clear that dissociated tumor cell culture models cannot faithfully replicate the spatiotemporal complexity of tumor biology.^9^ New technologies to probe intact tissues are needed to advance drug testing and personalized medicine. Over the past decade, investigators have increasingly utilized live tumor tissue to better replicate tumor physiology in pharmacodynamic and cancer biology experiments. Five approaches stand out: **1**) **tumor spheroids** (small spheres or “organoids” formed from patient-derived, dissociated cells),^10–17^ a model that can create cell-cell and cell-matrix 3-D interactions that more closely resemble in-vivo interactions and has been used for high-throughput drug screening assays that can be predictive of the patient’s responses,^10,11,18^ but retains only a limited amount of the original tumor microenvironment (TME); **2**) **micro-dissected tumors** based on sectioning of tumors into submillimeter tissue pieces that maintain the TME intact and are amenable to mass transport optimization and quantitative modeling;^19–25^ **3**) **tumor slices**, a technique often based on culturing thin slices of the tumor atop a porous membrane support^26,27^ and recently applied to cancer slices with great success,^28,29^ but sensitive to tissue scarcity; **4**) patient-derived xenograft (**PDX**) mouse models that permit study of drug responses in an intact organism, including immune checkpoint blockade in humanized PDX,^30^ however the rest of the TME is from the host mouse, and PDX from individual patients grow too slowly to inform initial post-operative therapeutic decisions; and **5**) **implantable or needle microdelivery devices**^31,32^ that locally deliver small doses of (up to 16) drugs to the tumor *in vivo*, with maximal preservation of the TME, but subject to limitations of tumor accessibility and patient safety.

Among these technologies, organotypic slice cultures have demonstrated the ability to closely resemble the TME architecture and allow for drug metabolism and toxicity studies in brain,^33^ liver,^34–36^ lung,^36,37^ kidney,^36,38^ and intestine^34^ tumor slices. Patient-derived clinical samples provide a unique tool to explore the susceptibility of individual tumors to specific therapies, but the small amount of tumor tissue extracted from any given patient limits the utility of this approach. We have developed a microfluidic platform for high-throughput chemosensitivity testing on individual slice cultures.^39,40^ Our platform has the potential to identify a subset of therapies of greatest potential to individual patients, on a timescale rapid enough to guide therapeutic decision-making.

Using our platform we have evaluated drug responses with U87 GBM xenograft tumors and showed similar and reproducible response profiles on and off device.^41^ This new method should allow for testing treatment sensitivity on the patient’s own tumor tissue to direct a physician’s therapeutic strategy. Additionally, our platform can provide a key missing link between drug testing in cell or animal models by testing new drug candidates in patient-derived cancer tissue in early stages of development.

Previously, we prototyped the platform by molding in polydimethylsiloxane (PDMS), i.e. by soft lithography.^39^ However, PDMS is not adequate for drug-based studies; both absorption into PDMS^42^ and adsorption onto PDMS^43^ can potentially alter experimental outcomes by changing the target concentrations and by partitioning molecules in undesired regions of a microfluidic device. PMMA features less drug absorption than PDMS, although it can still adsorb drugs on its surface.^44^ Furthermore, our early PDMS prototype involved more than two days of highly-skilled fabrication per unit, so it was not readily translatable for large-scale fabrication for widespread adoption. We explored the 3D-printing of PDMS as a possible manufacturing solution,^45^ but the printed elastomer is virtually identical to Sylgard 184, so it does not address the drug binding problems; we have also demonstrated high-resolution 3D-printing in inert biocompatible resins,^46^ but limitations in the build size of commercial 3D-printers prevented its use in this application.

Here we report a thermoplastic version of the platform made entirely in poly (methyl methacrylate) (PMMA) by digital manufacturing based on CO_2_ laser micromachining. PMMA has been widely used to build microfluidic devices for biomedical applications because it is biocompatible, has excellent optical transparency, low gas permeability, and is less expensive than PDMS.^47–49^ Compared to conventional photolithography and soft lithography methods, digital manufacturing shortens the design and processing time, streamlines assembly, and reduces manufacturing costs. This manuscript reports a detailed description of the complete fabrication process including CO_2_ laser optimization for microfabrication, post-processing, and solvent bonding techniques for assembly, as well as measurements of diffusion between and above channels using the device. We demonstrate the functionality of the platform by the multiplexed delivery of anti-cancer drugs onto glioblastoma multiforme (GBM) xenografts and onto patient-derived colorectal cancer (CRC) tumor slices. We further demonstrate the versatility of the device with an expanded drug treatment panel, multiple types of fluorescent cell death indicators, and real-time measurements using the device.

## Experimental

### CO_2_ Laser Micromachining

The current version of our platform consists of a 40-well plate with an integrated channel network layer. We fabricated the platform by laser micromachining of PMMA substrates and thermal fusion and solvent bonding techniques. The device is composed of three layers: a 19 mm-thick PMMA bottomless 40-well “well plate” (1227T569, McMaster-Carr, Elmhurst, IL), a 300 μm-thick PMMA channel network layer (Astra Products, Baldwin, NY (11510103)) containing 40 closed microchannels that feed into 40 roofless microchannels (“delivery channels”), and a 125 μm-thick PMMA sealing layer (AFT00, SPolytech, Chungbuk, Korea). In addition to the main components, the device also has a customized base and a lid (1227T569, McMaster-Carr, Elmhurst, IL). The base raises the device from the surface to avoid scratches (thus maintaining optical clarity) and makes its dimensions compatible with conventional imaging stages. The lid prevents contamination and allows proper air flow for tissue culture. During operation, the roofless delivery channels are sealed with a porous membrane (see below).

The CO_2_ laser system used (VLS3.60, Scottsdale, USA) has a wavelength of 10.6 μm and a maximum power of 30 W. We optimized the power and speed settings of the CO_2_ laser to achieve specific widths and depths for the microchannels and to cut the outlines of the channel network and sealing layers. To develop the well plate, we utilized another CO_2_ laser system (ILS12.150D, Scottsdale, USA), which has a maximum power of 150 W and the same wavelength. Multiple passes with this laser were needed to create smooth through-cuts for the wells of the well plate. We also cut out a hole in the plate to form the main outlet of the device. Finally, we engraved a shallow well around the outlet hole for tubing installation after assembly.

### Post-ablation processing

Laser ablation of PMMA includes both polymer debris and reflow. To remove debris from the laser-cut substrates, we rinsed each of the device components with DI water and sonicated them in an IPA bath for 2 min. To reduce surface roughness and improve the optical quality, we exposed the channel network to chloroform vapor. We used a glass container (264 mm (L) × 213 mm (W) × 165 (T) mm) filled with 50 mL of chloroform and steel standoffs (6 mm) to elevate the laser-micromachined layers 3 mm above the chloroform surface. We exposed the channel network layer and the sealing layer to chloroform for 4 and 2 min, respectively.

Laser cutting of the well-plate caused the PMMA substrate to warp (∼1 mm warpage over 113 mm length) due to the thermal stresses induced by the laser during ablation. Assuming the PMMA substrate was placed in a vacuum chamber, with no heat sink, and uniform heat distribution, it would absorb about 135,000 J of energy after laser cutting at 150W for 15 min. Considering that the laser platform has a heat sink (air assist and an aluminum pin table) and there is abundant heat loss due to a) reflected longwave radiation, b) re-radiated longwave radiation, and c) heat transmission, the actual heat absorbed by the PMMA plate should be much lower. Nevertheless, we observed that the prolonged laser ablation processes caused melting and deformation of the PMMA well plate. We were able to fix the deformation of the plate after ablation by pressing at 1,000-1,500 psi and heating at 110 °C at the same time for 10 min in a thermal press (Carver Inc. 4126). Afterwards, the plate was cooled down to room temperature for 10 min while still being pressed. The corrected well plate was then rinsed with DI water, sonicated with IPA, and exposed to the chloroform vapor for 30 min using the previously mentioned setup.

### Thermal fusion and solvent bonding

Exposure to chloroform vapor also causes the PMMA to become slightly adhesive by inducing polymer reflow.^50,51^ After chloroform vapor treatment, the surface of the PMMA substrates becomes soft due to polymer solvation. When two treated surfaces are placed in contact with each other, a cohesive molecular bond is formed while excess vapor evaporates from the interface. For assembly, we exposed the channel network layer and the sealing layer to chloroform vapor. For thermal fusion bonding, we first hand pressed the sealing layer onto the channel network layer to form a weak bond. Then, to ensure uniform bonding, we sandwiched the two layers between two ∼3 mm-thick PDMS slabs with the same outer dimensions as the channel network layer. Finally, we placed the whole ensemble in the heat press for 10 min between 120-160 psi at 60°C.

Although it is possible to build complex microfluidic networks with laser-cut laminates by using glue as the bonding layer,^52^ biocompatibility concerns about the glue prompted us to adopt a glue-free approach based on solvent-bonding. To bond the channel network layer to the 40-well plate we used methylene chloride (Weld-On 4, Durham, USA). This solvent also softens the surfaces of the PMMA substrates, and it bonds the substrates together as it evaporates. For this bonding process, we directly exposed the bottom of the plate to methylene chloride for 15 seconds. Immediately after, we blew nitrogen into the outlet tubing hole of the 40 well-plate to remove excess solvent; if left behind, this excess solvent can dissolve the channel inlets and outlet in the channel network layer during the bonding process. Then, we aligned the channel network layer with the outlet of the well-plate and hand-pressed them together. We placed a 500 μm-thick blank PMMA sheet with the same outer dimensions as the channel network layer on top of the channel network layer, and then we put all the layers in between two PDMS slabs for even pressure distribution. We then placed the whole assembly in the heat press at 200 psi for 5 min at room temperature. To build the platform prototype that can accept Transwells, we followed the same pressing process and we bonded the device with 3M™300 LSE transfer adhesive. We first bonded a set of two laser-cut 1/8” PMMA well-plates with the modified design followed by the channel layer. Finally, we added the connection tubing to the outlet in the well-plate and secured it into place by filling the empty well surrounding the outlet with ethyl cyanoacrylate glue (Gorilla Glue, Cincinnati, USA).

### Hydrophilization and sterilization

After device assembly and prior to use, we treated each device with oxygen plasma for 5 min at 660 mTorr to increase the hydrophilicity of the PMMA surfaces. Then, to prepare the device for use with biological samples, we placed the device in a tissue culture hood and filled the microchannels by pipetting sterile PBS into the well reservoirs and the central culture area. We covered the roofless channels with a PDMS membrane and suction was applied to completely fill the microchannels. Once the microchannels were filled, we left the device under UV for 1 hr for sterilization.

### Device operation

Before use, we filled the device with culture medium and transferred it to a cell culture incubator to allow the temperature and pH to equilibrate. After ∼1 hr of incubation, we transferred the slices from the tissue culture insert by cutting out the PTFE membrane and placing it onto the roofless channels of the device. After transferring the slices to the device, we imaged the central culture area to capture position of the tissue slices relative to the delivery channels. Then, we filled each well reservoir with either drug or buffer with at least one buffer lane between each drug delivery channel. We diluted drugs (MedChem Express) from DMSO stocks (10-200 mM), except for cisplatin (3M stock in dH_2_O). We operated the device by connecting the outlet of the device to a 60 mL syringe (BD Bioscience, San Jose, CA) and syringe pump (Fusion 200, Chemyx Inc., Stafford, TX) at a flow rate of 1.5 mL/hr for xenograft drug studies and 2 mL/hr for vertical diffusion and CRC studies.

### Scanning Electron Microscopy

We took scanning electron microscopy (SEM) images of the laser-cut PMMA channels to evaluate topological changes of the channels due to chloroform exposure and the dimensions of the channels (roofless and sealed). We prepared the samples by rinsing with DI water, sonicating with IPA for 2 min, and rinsing with DI water again. We utilized nitrogen gas to remove excess water and left the samples dry overnight. To image microchannel cross-sections, we prepared substrate samples through a freeze-fracture process involving immersion in liquid nitrogen. Finally, prior to SEM observation, we coated all samples with a thin, 19.5 nm-thick film of Au-Pd to prevent charging.

### Image acquisition

We imaged the tissue slices for epifluorescence and brightfield imaging using a Nikon Eclipse Ti inverted microscope (Nikon Instruments, Melville, NY), except for where specified in Fig. 7l where we used a Nikon A1R Confocal system (Lynn & Mike Garvey Imaging Core, University of Washington). We acquired images using an automated XY stage and 4×, 10×, and 20× objectives. To generate images, we stitched all the images with 10% overlap. For controlled imaged acquisition, we used the Nikon NIS-Elements AR software.

**Figure 1.**
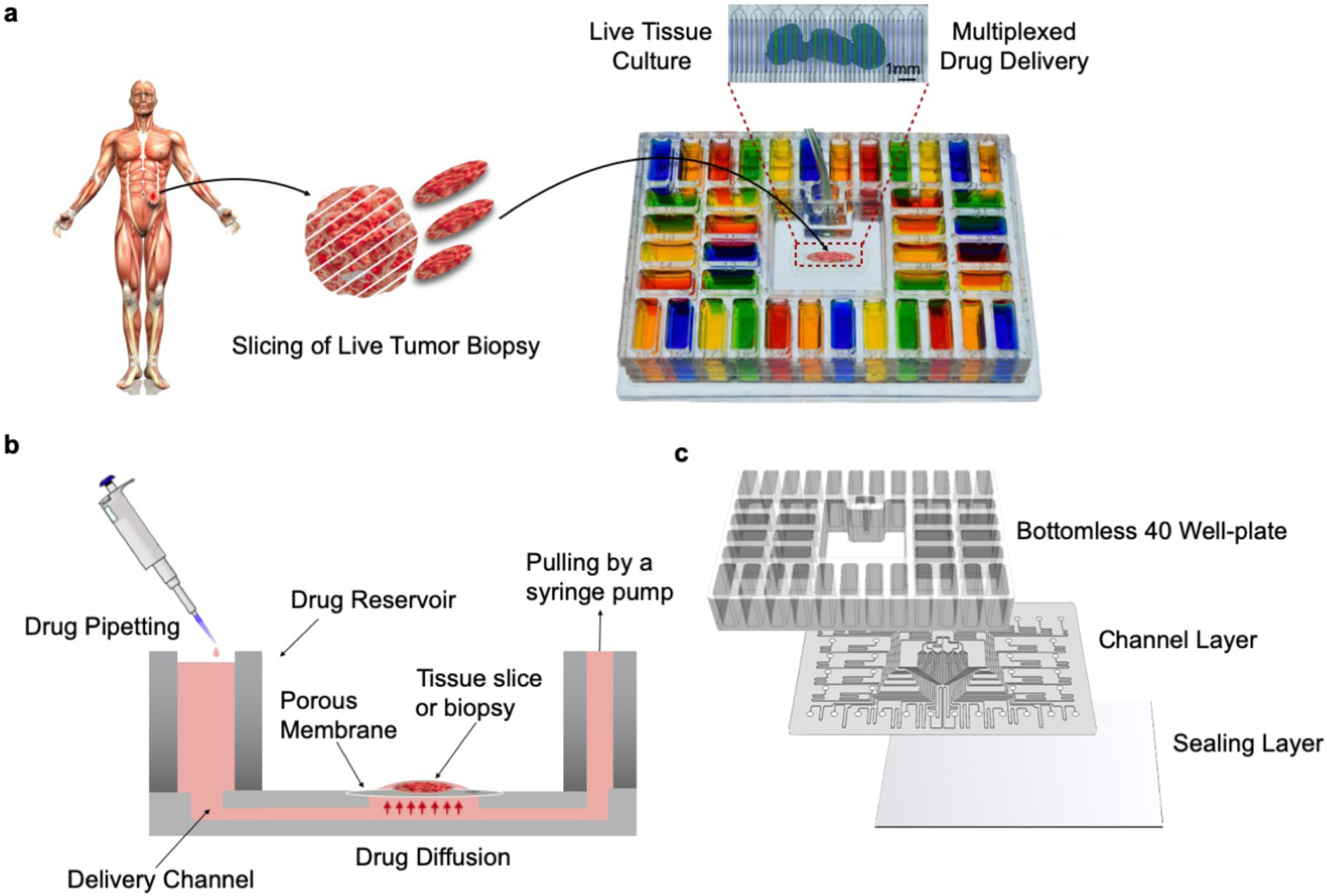
Microfluidic device design and overview. (a) Proposed application and device functionality. Micrographs of a mouse glioma tumor slice exposed to two different cell nuclear binding agents (Hoechst, blue, and Sytox Green, green) through alternating streams. (b) Cross-sectional schematic of the device. The device is operated by gravity flow and the total flow rate is driven by a syringe pump through a common outlet: one syringe pump controls flow across all 40 fluidic streams. Tissue slices are cultured on a PTFE porous membrane. The wet membrane seals the roofless microchannels by capillarity, which allows for fluidic stream transport of culture medium to tissue. (c) Exploded schematic of the PMMA platform showing from top to bottom: 1) bottomless plate with 40 inlet wells, 2) 300*µ*m-thick channel network layer, and 3) 125 *µ*m-thick sealing layer.

**Figure 2.**
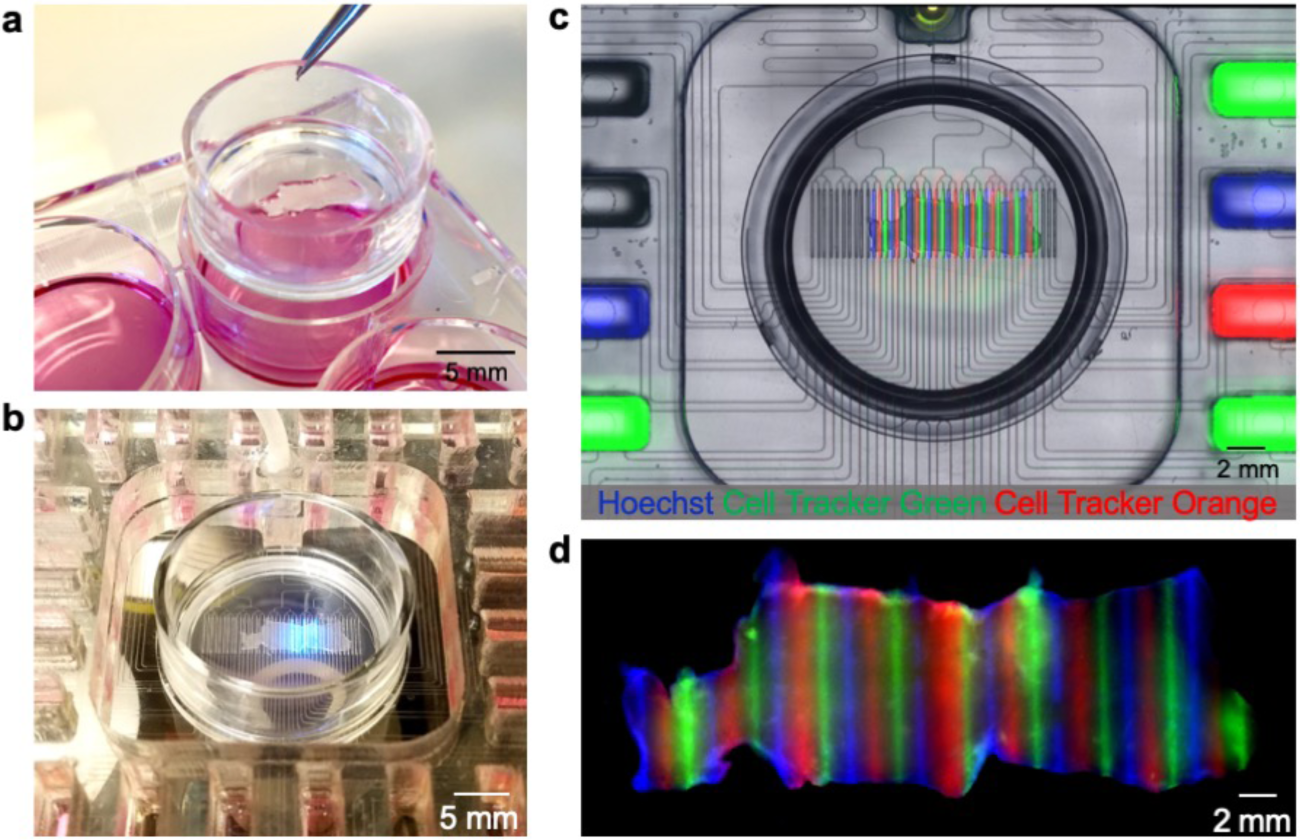
Microfluidic device with Transwell insert. (a) Photograph of standard six-well plate with Transwell insert containing U87 GBM flank xenograft slices. (b) Live imaging with the platform with incorporated Transwell insert. (c) Fluorescent image showing microfluidic delivery of Hoechst (blue), Cell Tracker Green (green), and Cell Tracker Orange (red) to live GBM slices. (d) Fluorescent image showing fixed tissue 48-hours after delivery.

**Figure 3.**
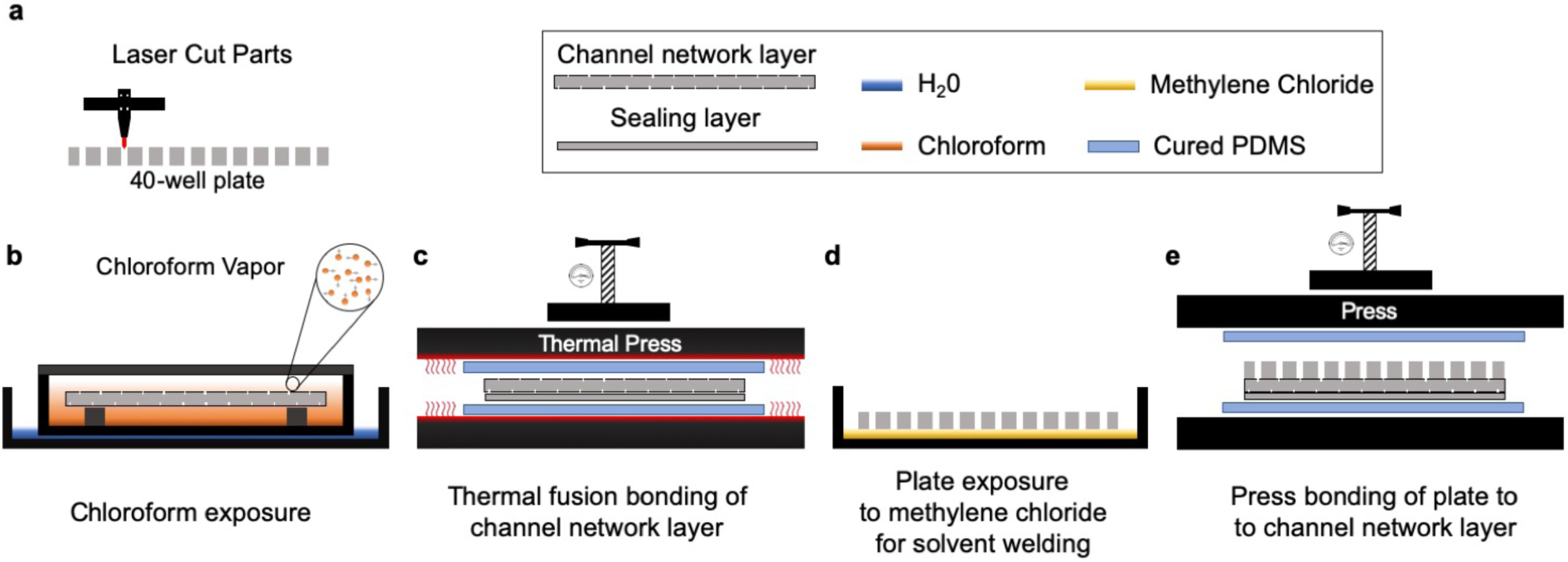
General overview of device fabrication protocol. The overall fabrication of the device consists of (a) laser cutting/engraving and cleaning each of the laminates that compose the microfluidic platform (30 min); (b) chloroform vapor exposure (6 min); (c) thermal fusion bonding (5 min); (d) solvation of the under surface of the 40-well plate with methylene chloride (1 min); (e) alignment and assembly by pressing (5 min); and outlet/base installation (10 min, step not shown).

**Figure 4.**
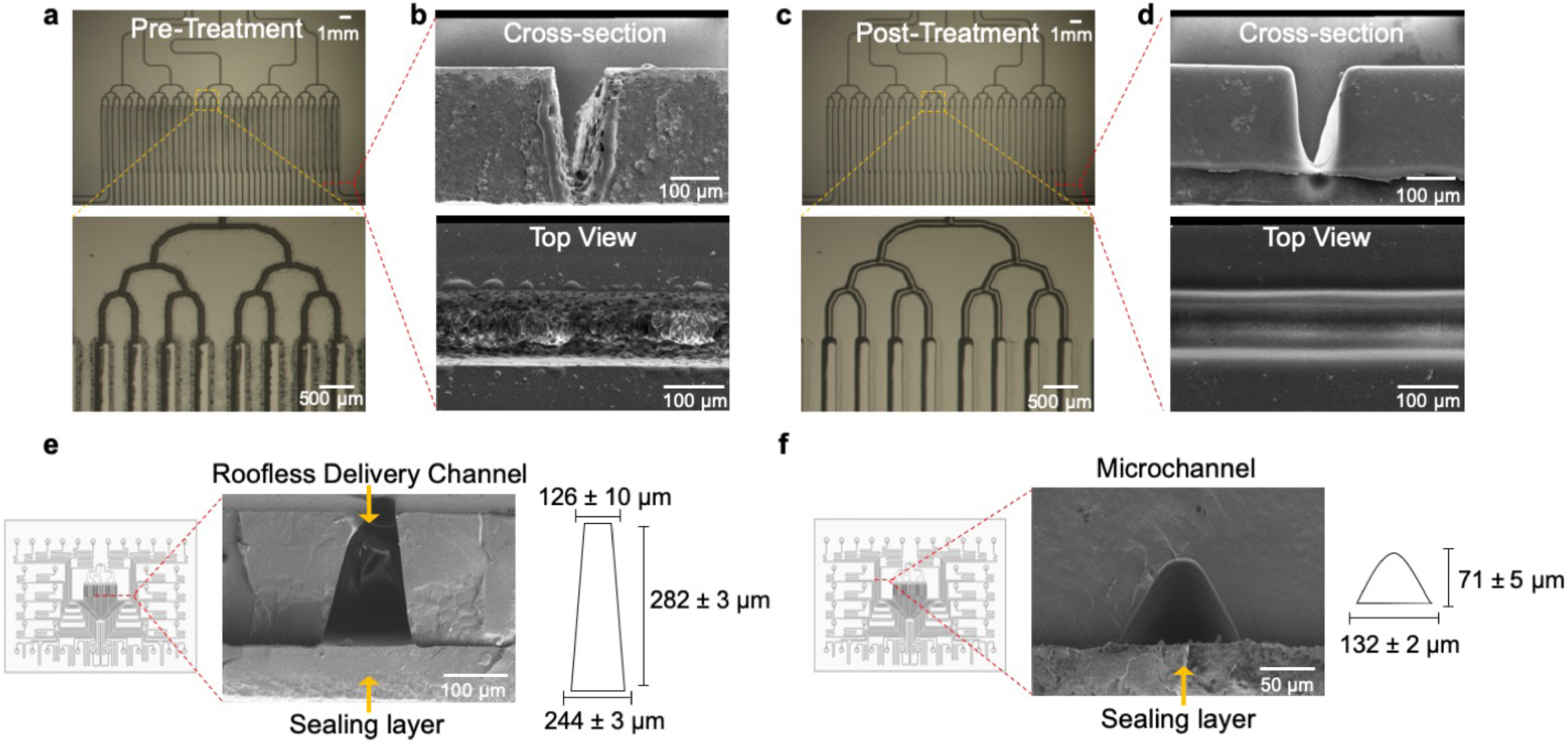
Smoothing of channel features by chloroform exposure. (a) Micrographs of the channel network layer prior to exposure to chloroform vapor. The channels have poor optical clarity as evidenced by the darkness of the channels. (b) SEM images of the cross-sectional view and top view of channels after being engraved by laser micromachining. The channels have a rough, porous structure and bulges around the channel rims. (c) Micrographs of the channel network layer after exposure to chloroform vapor. The channels have increased optical clarity since more light penetrates through the channels. (d) SEM images of the cross-sectional view and top view of the channels after being exposed to chloroform vapor. The channel profile becomes smoother after exposure to chloroform and the surface roughness is reduced. (e) SEM of the cross-sectional view of the roofless/delivery microchannels. The roofless delivery channels have a trapezoidal shape where the opening facing the sealing layer is 244 ± 3 μm wide, the opening facing the membrane and tissue sample is 126 ± 10 μm wide, and the depth is 282 ± 3 μm. (f) SEM of a sealed microchannel. Solvent bonding with chloroform vapor exposure and thermal pressing yields sealed microchannels. The laser micromachined channels have a curved shape with a width of 132 ± 2 μm and a depth of 71 ± 5 μm.

**Figure 5.**
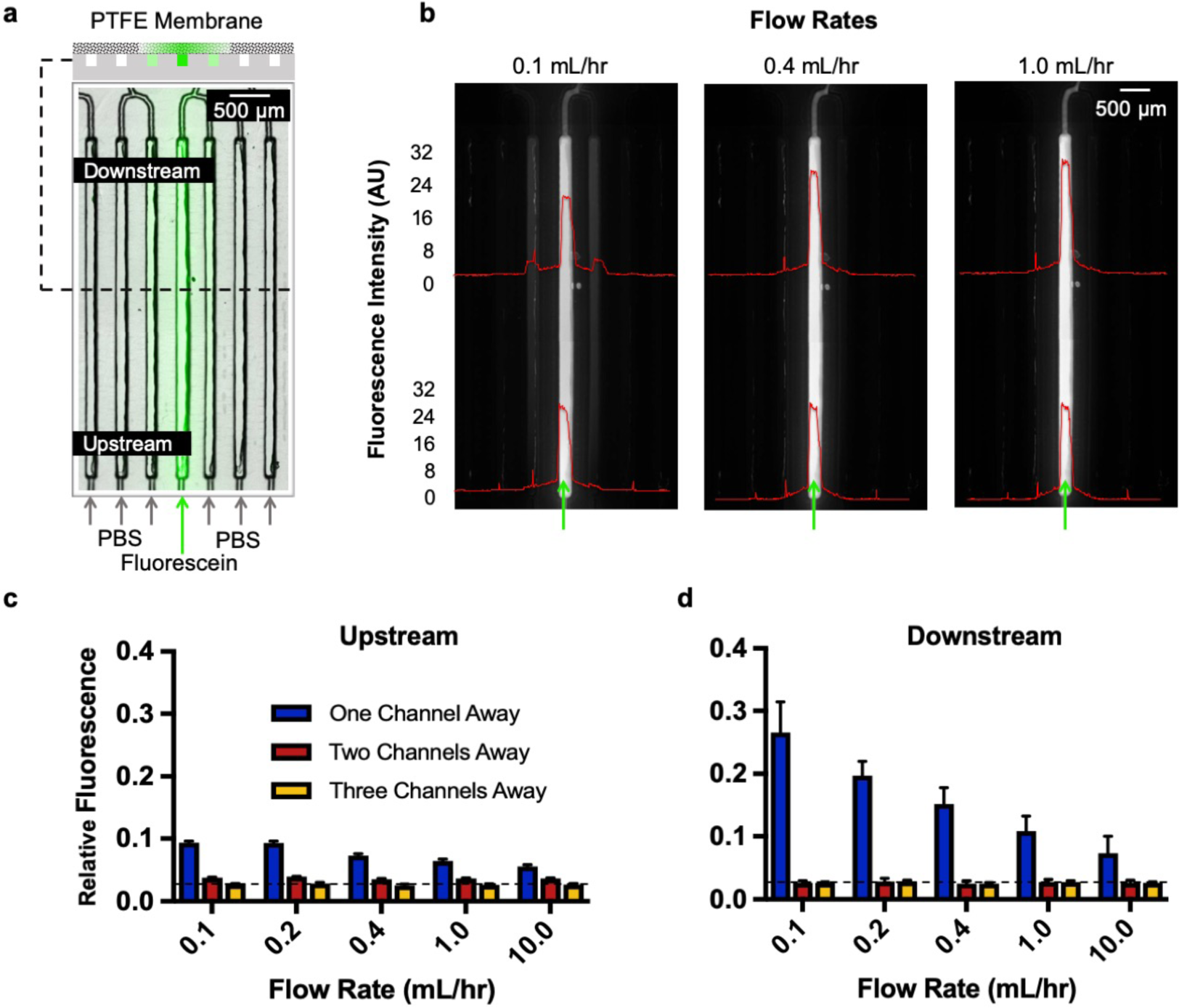
Lateral diffusion assessment using fluorescein. (a) Delivery schematic of 100 mM fluorescein across 6 of the 40 delivery channels, with the rest having PBS. We intentionally isolated each of the 6 delivery channels by 6 channels to analyze spread three channels over on both directions when possible as shown in the top cross-section schematic. (b) Fluorescent images taken at 0.1, 0.4, and 1 mL/hr showing overall diffusion profiles upstream and downstream of the delivery channels and its adjoining channels. (c) Upstream and (d) downstream measure of the relative fluorescence showing lateral spread one, two, and three channels away from the delivery channel.

**Figure 6.**
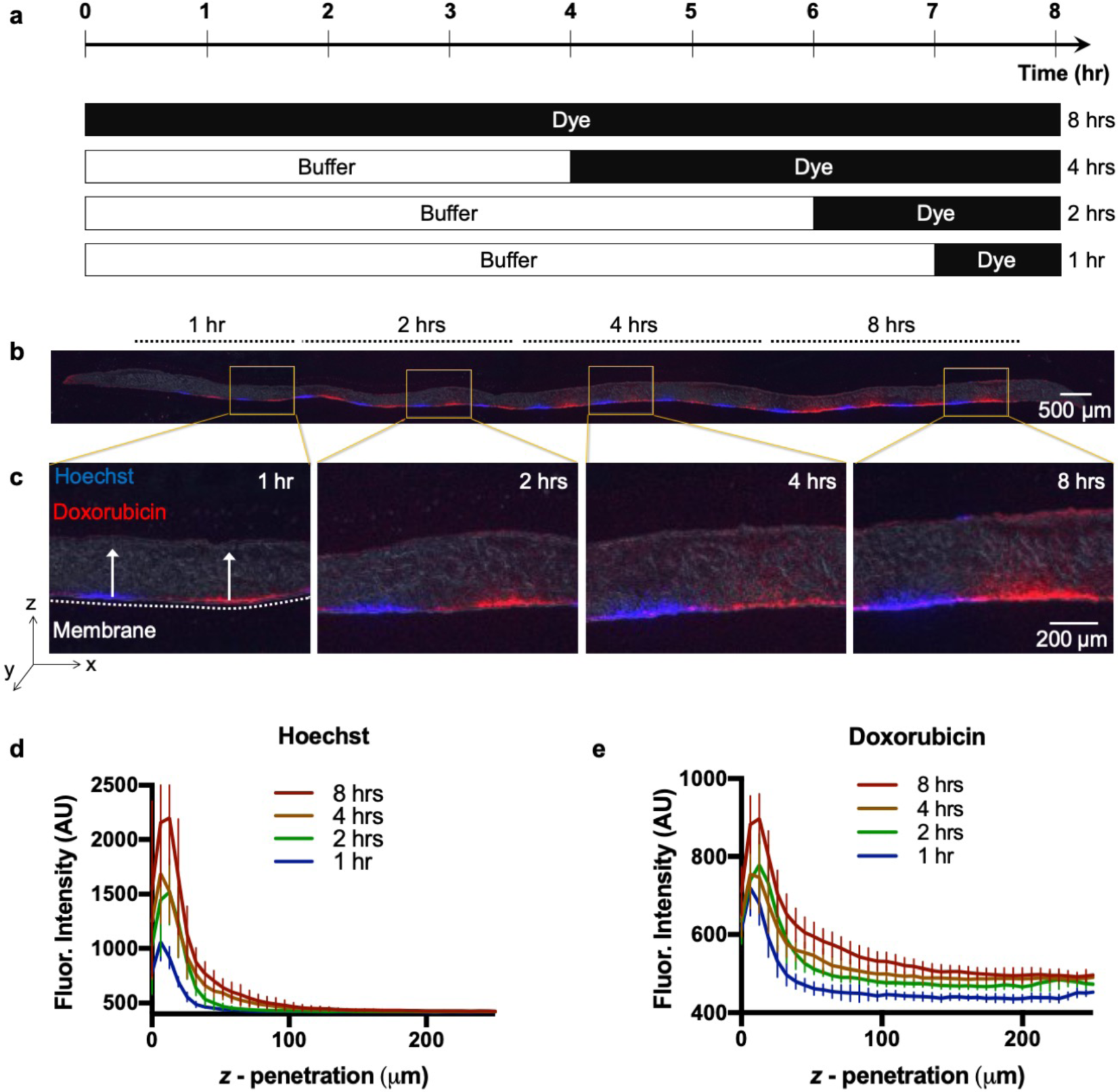
Vertical diffusion into U87 xenograft tumor slice. a) Schematic of timed delivery of Hoechst and Doxorubicin. b) U87 tissue section (10 μm-thick) after 1, 2, 4, and 8 hrs Hoechst (blue) and Doxorubicin (red) exposure. c) Section magnification of exposed areas. d) Intensity profiles showing Hoechst. (e) Doxorubicin tissue z-penetration for each time point (n=8, per time point). Ave ± SD.

**Figure 7.**
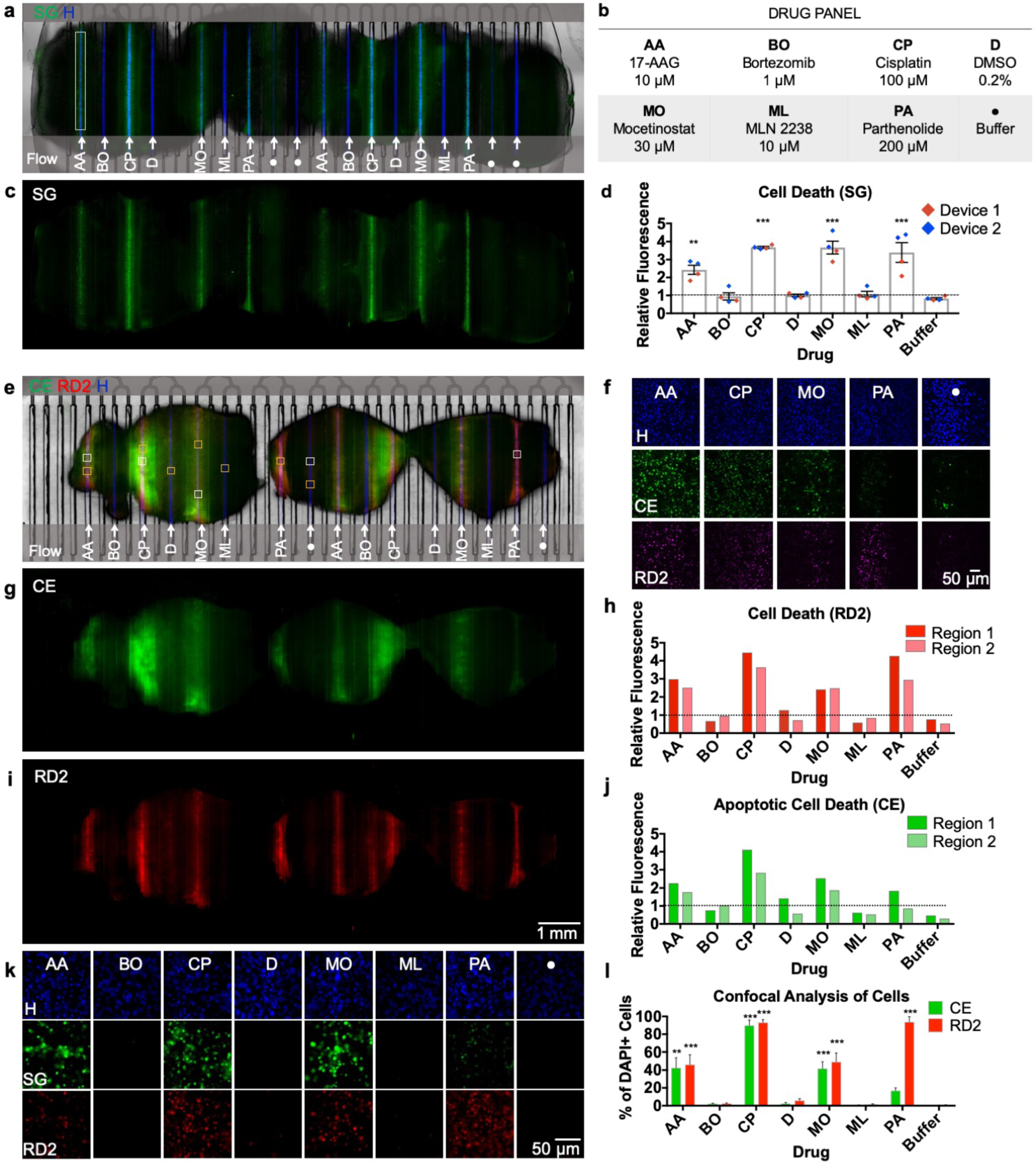
Multiplexed drug exposure of U87 GBM xenograft flank tumor slices. a) On-device micrograph of U87 flank xenograft slices (3) after 48-hour drug exposure showing selective delivery and response to 8 different conditions (b). Hoechst nuclear dye (blue) denotes areas of delivery and Sytox Green areas of cellular death (c). d) Drug response measured by relative fluorescence (to DMSO) obtained from two different devices ran in parallel. e) On-device micrograph of U87 flank xenograft slices (3) after 48-hour drug exposure, labeled with Cell Event (green, g) and RedDot 2 (red, i). f) Confocal micrographs (20×) of drugs showing response compared to a negative control (samples taken from white boxes in (e)). h) Apoptotic response shown by Cell Event from two different tissue regions (dotted line = DMSO vehicle control baseline). j) Necrotic and late apoptotic areas shown by RedDot 2 from two different tissue regions (dotted line = DMSO vehicle control baseline). k) 20× fluorescent micrographs for each condition (yellow boxes in (e)). l) Cell Profiler cellular analysis showing % of DAPI+ cells positive for Cell Event/RedDot 2. n = 11 (AA), 5(BO & CP), 8(D & ML), 7(MO & Buffer), 6 (PA). Ave ± SEM. One-way ANOVA versus DMSO with Dunnett’s multiple comparison test. *p<0.05, **p<0.01, ***p<0.001. Abbreviations: CE (Cell Event), SG (Sytox Green), RD2 (RedDot 2), H (Hoechst).

### Lateral spread assessment using fluorescein

We covered the roofless delivery channel area (19.70 mm × 5.42 mm) with a manually-cut porous Millicell® polytetrafluoroethylene (PTFE) membrane (32 mm × 20 mm, 0.4 μm, Millipore). We filled the well reservoirs of 6 channels with 100 μM fluorescein, leaving the rest filled with phosphate-buffered saline (PBS). To analyze spread three channels in both directions (when possible), we intentionally separated each of the selected delivery channels by 6 channels. After loading the device, we operated it at 0.1, 0.2, 0.3, 0.4, 0.75, 1, and 10 mL/hr. To determine the fluorescence profiles of fluorescein, we took images at each flow rate after waiting for equilibration for ∼10 min. We also acquired images at 7 ms and 70 ms exposures for every flow rate. Then, we analyzed lateral diffusion in the middle, upstream, and downstream, for each of the 6 channels of interest and the adjoining channels. In our first analysis, we did fluorescence profile along the delivery channels and its three adjacent channels. In the second analysis, we measured the relative fluorescence in each of the three channels adjacent to the delivery channel after background subtraction. For each flow rate, we averaged the values based on the location of the channel: 7 ms for the delivery channel and 70 ms for one, two, or three channels away. We utilized relative fluorescence between the two exposures to obtain a correction factor to adjust the 7 ms data to match the 70 ms data. Finally, we normalized the data to the average fluorescence at the 6 delivery channels.

### Diffusion of fluorescent compounds in live tissue

After 7 days in culture, we transferred the U87 flank xenograft slices to the device, then exposed them to alternating delivery channels containing Hoechst (16 μM) and Doxorubicin (DOX, 10 μM) for four different time periods (1, 2, 4, and 8 hrs). There were 2 lanes for 1 hr and 4 lanes for 2, 4, and 8 hrs. After exposure, we rinsed the delivery channels with PBS for 10 min. To prevent any additional diffusion, we then immediately froze the exposed slices and analyzed as dry cryosections (10 μm thick). For vertical diffusion analysis, we placed vertical profiles (100 μm wide) in the center of each delivery location. We averaged three adjacent sections to yield profiles for each of 8 locations per time period (2 locations on each delivery lane). We plotted the average and standard deviation (SD), with n=8 (except for n=4 for 1 hr).

### Diffusion constant estimation

We applied Fick’s laws of diffusion to quantitatively understand the penetration of the drug within the tissue. A solution to the Fick’s second law of diffusion in semi-infinite media and a constant concentration surface is given by:

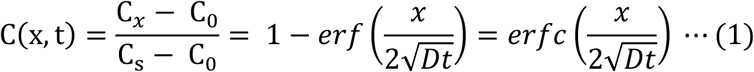

where *D* represents the effective diffusivity of Hoechst and doxorubicin in tissue, C(x = 0) = C_S_ is the concentration of doxorubicin at the PTFE membrane and tissue interface and C(x = ∞) = C_0_ corresponds to the initial concentration of doxorubicin on the top surface of the tissue. We assume that C_S_ remains constant over time, and C_0_ = 0 for the early time-periods (< 8 hrs) of drug application (that is, the drug has not traversed the entire thickness of the tissue). The characteristic diffusion length (L) at a given time (t) is defined as the distance at which the concentration of the diffusing species reaches ∼50% of the source concentration (C_S_) and can be approximated by 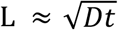. We determined the diffusivity (D) of Hoechst and doxorubicin in the tissue from the experimentally observed diffusion length (L) at each time point (t).

### GBM xenograft slice culture

We cultured U-87 MG cells (U87, ATCC) in DMEM/F12 (Invitrogen) supplemented with 10% fetal bovine serum and penicillin/streptomycin. We passaged the cells every 3-5 days at ∼75% confluency. Mice were handled in accordance with a protocol approved by the University of Washington Animal Care and Use Committee. We injected male immunodeficient nude mice (Taconic, Foxn1^nu^) aged 4-10 weeks subcutaneously in the flank (∼1 million cells in 200 μL of serum and antibiotic free medium). Before the flank tumor volume reached 2 cm (2-4 weeks), we sacrificed the mice. Once sacrificed, we removed the tumor, cut 250 μm-thick slices with a 5100mz vibratome (Lafayette Instrument), and cultured on top of PTFE, 0.4 μm pore membrane Millicell® cell culture inserts (Millipore) in 6-well plates. The slice culture medium underneath contained Neurobasal-A medium (Invitrogen) with 25% heat-inactivated horse serum (Sigma), Glutamax (Invitrogen), 2× penicillin/streptomycin (Invitrogen), and growth factors (EGF 20 ng/mL and FGF 20 ng/mL, Preprotech or Invitrogen). We changed culture medium three times per week.

### Patient-derived tumor slices

We obtained metastatic rectal cancer tumor tissue with informed consent and treated in accordance with Institutional Review Board approved protocols at the University of Washington, Seattle. We prepared CRC tumor slices from a 68-year-old male with metastatic rectal cancer post-neoadjuvant chemotherapy, resection, and radiation. Slices were treated with two different standard chemotherapy combinations for metastatic CRC: 1) Fluorouacil (5FU) and Oxaliplatin, 1 μg/mL each (termed “FOLFOX”), and 2) 5FU and Irinotecan, 1 μg/mL each (termed “FOLFIRI”). Prior to tumor resection, the patient had FOLFOX treatments for about 7 months (cycles sometimes excluded Oxaliplatin due to side effects). After chemotherapy, the tumor size was ∼3.7 cm. We sectioned the CRC tumor from 6 mm core punches taken from the resected tumor. With the 6 mm cores, we cut 250 μm-thick slices with a vibratome (Leica) and placed them in culture with shaking. We have found that CRC slices maintain their histology, viability, and proliferation for up to 1 week in culture.^53^ After 3 days of culture, we quantified change in viability after drug exposure using CellTiter 96® Non-Radioactive Cell Proliferation Assay (MTT, Promega). We performed the MTT assays in a 48 well plate containing 400 *µ*L culture media and 80 *µ*L of MTS reagent in each well. After transferring the slices to each well with a sterile pipette tip, we placed the slices in a rocker inside the incubator for 3 hrs. After 3 hrs of incubation, we placed 200 *µ*L from each well into a 96 well plate and read absorbance at 490 nm. For off-device analyses, we determined response through a viability comparison before and after treatment using fluorescence microscopy.

### Multidrug exposure data analysis

We used the free Fiji^54^ image analysis program for SG and CellEvent/RedDot2 image analysis on tiled 2× images taken with tumor slices on the microfluidic platform. For the device experiments, we selected a rectangular region (∼130 μm wide) for each condition using the Hoechst channel to avoid ∼25 μm of the edge of the slices (see Fig. 6a). For background subtraction on both conditions, we used four circular regions outside the slices. We calculated the background by averaging all values for each fluorescent channel. After background subtraction, we calculated the total average fluorescence for each region relative to that of DMSO (here termed “vehicle control” because the drugs were solubilized in DMSO; all solutions contained 0.1% DMSO independently of the drug concentration except for cisplatin). We analyzed the images acquired by confocal microscopy (20×) with CellProfiler (Broad Institute). We created a custom routine to identify all nuclei from each channel (Hoechst, CellEvent, RedDot2) using a single manual threshold applied to all images for each channel.

For CRC experiments, we took 20× images of DAB staining for CC3 at all areas of drug exposure from seven sections of the slice cut perpendicular to the membrane. To remove cell counting bias, the observer that manually counted CC3+ or Ki67+ cells only in tumor+ areas was blinded to the experimental conditions (except for off-device experiments). For automated analysis of CC3+ or Ki67+ area, Fiji was used to manually remove non-tumor areas (Ki67), to perform background subtraction, to separate the brown staining (color deconvolution plug-in with “H DAB”), and to determine a single manual threshold for positive staining for all of the conditions. For Ki67 experiments, we took 10× images covering each entire section, and used Fiji to divide the images into 100 μm wide images aligned on the center of treatment conditions. Then Cell Profiler^55^ was used to determine the overall tissue area, tumor area, and positive staining area, using a previously determined single manual threshold for all images. The 200 μm-wide regions centered on each treatment area were used for each condition.

### Live-tissue staining and post-tissue processing

For the experiments where we used the Transwell version of the device, we used Hoechst (Invitrogen, 16 μM), Cell Tracker Green CMFDA (Invitrogen, 10 μM) and Cell Tracker Orange CMRA (Invitrogen, 10 μM). After transferring the tissue culture insert containing slices, we aspirated growth medium from the wells and added each of the fluorescent dyes in an alternating order and ran the device at 1.5 mL/hr for 2 hrs.

For drug exposure studies, we aspirated the drug from the well reservoirs 2 hrs before the end of the experiments. Subsequently, we filled the empty drug reservoirs with growth medium containing Hoechst (Invitrogen, 16 μM) and SYTOX green (SG, Invitrogen 0.01 μM), or with CellEvent (1/1000, Invitrogen) and RedDot 2 (1/400, Biotium), to label the areas that had drug exposure and to assay cell death. After exposure to these solutions for 2 hrs, we changed the medium in all the well reservoirs to PBS for a 30 min wash. After on-device imaging, we transferred the slices to a 6-well plate, fixed with 4% paraformaldehyde overnight, and then cryoprotected with 30% sucrose/PBS overnight two times.

Similarly, for the CRC device experiments, we exposed the slices to Hoechst in the drug delivery channels for the last 2 hrs on device, then transferred to a 6-well plate and exposed them to CellEvent (1/1000, Invitrogen) for 1 hr. After fixation and cryoprotection as above, we cut the slices in half, perpendicular to the delivery channels, and processed them for cryosectioning (10 μm thickness). For immunostaining, we pretreated tissue sections with 0.6% hydrogen peroxide in methanol for 30 min, washed, and then processed for antigen retrieval by steaming for 30 min in 10 mM sodium citrate, 0.05% Tween 20 (Sigma), pH 6.0. After at least 30-min incubation in blocking solution (Tris-NaCl-blocking buffer or TNB buffer, Perkin Elmer, with 0.1% Triton X-100), we incubated the tissues with rabbit anti-active cleaved caspase 3 (CC3, 1/600, Cell Signaling) or Ki-67 (1/1,000, AbCAM, ab15580) primary antibodies (diluted in TNB) overnight at 4°C. Finally, we incubated the tissues with peroxidase polymers of the appropriate species for 30 min (rabbit from Vector Labs MP7401 or mouse from Biocare MM510) then with the chromogen 3,3’-Diaminobenzidine (DAB, Vector Labs) and lightly counterstained with hematoxylin. We performed all tissue washes with PBS.

For off-device CRC experiments, we embedded fixed slices in paraffin, cut 4 *µ*m-thick sections and placed them on glass slides. After deparaffinization, rehydration, and quenching, we conducted antigen unmasking in 0.01 M citrate buffer of pH 6.0 in a microwave for 10 min. Subsequently, we incubated the slides with CC3 (1/200, Cell Signaling) and Ki67 (1/200, Dako) primary antibodies. After incubation, we developed the sections with an avidin–biotin technique using the VECTASTAIN Elite ABC kit (Vector Laboratories, Burlingame, CA). Finally, we counter stained the slides with Hematoxylin QS and mounted with Permount (Fisher Scientific, Santa Clara, CA).

## Results and Discussion

### Microfluidic device design

Our patented^40^ platform permits regioselective delivery of drugs with spatiotemporal control on live tissue slice cultures (Fig. 1a, b). The platform consists of three functional components, as well as a lid and a base. From top to bottom these components are: 1) a 40-well plate with a central culture area, 2) a microfluidic channel network layer, and 3) a sealing layer for the bottom surface of the channel network (Fig. 1c). We fabricated the device using a combination of CO_2_ laser micromachining and solvent bonding techniques. After optimizing both techniques, we irreversibly bonded all the PMMA components to produce a leak-proof platform.

The microfluidic platform consists of 40 microchannels connecting to a central tissue culture area for drug delivery. Upstream of the central area, each microchannel is individually connected to a different reservoir well of the 40 well-plate, containing a drug. Downstream of the delivery area, all the microchannels are connected to a common outlet via binaries. The delivery microchannels, initially “roofless”, are separated by 250 μm-thick PMMA walls, designed to sit underneath a PTFE porous membrane. When a wet PTFE porous membrane is placed onto the roofless microchannels, capillary forces cause the membrane to adhere to the surface; the membrane then becomes the “roof” for the roofless microchannels. (We note that it is possible to generate flow in roofless microchannels by application of positive pressure at the inlet,^56,57^ but here we wished to generate flow by suction through the outlet, so adding a roof provides a zero-flow constraint and forces flow from the inlets.) Application of negative pressure at the outlet causes a slow flow under the membrane, increases the PTFE membrane adhesion to the PMMA surface to “seal” the channel, and allows for the diffusive supply of different reagents to the tissue in different channels. Equal microchannel resistances, achieved by having equal microchannel lengths and curves across the whole architecture, result in equal flow across microchannels (see ESI. Fig. S1 and Flow Characterization section below). A constant CO_2_ laser power and speed creates microchannels with uniform width and depth, although small imperfections in the channels occur due to the stochastic nature of laser melting and tracking errors in the mechanisms that move the laser.

The platform allows for live tissue culture and high-density delivery of up to 40 different solutions. Live tumor slices are cultured on top of the porous membrane that forms the roof of the open microchannels in the central culture area of the device. Each microchannel is fed by a 1.6 mL reservoir. With a flow rate of 1 mL/hr, each well could provide up to ∼64 hr of uninterrupted reagent delivery. To operate the device, the user fills the reservoirs of the 40 well-plate and activates flow with a syringe pump connected to the outlet of the device. Solutions move through the channels on the bottom surface and then diffuse upwards through the membrane into the tissue (Fig. 1b). The platform is easy to transport, and the clear plastic and well-plate footprint make the device compatible with standard imaging systems.

The interfacing of microfluidics with Transwell inserts has allowed for the manipulation of the basal surface of cells in culture^58^ and represents a very intuitive approach to cell-based microfluidics. Our device can also be adapted to facilitate the transfer of cultured tissue onto the device from Transwell inserts (Fig. 2a), which are made of the same porous membrane that we use for the platform. After manually removing the ∼2 mm feet at the bottom of the insert, the entire Transwell insert can be directly placed into the device (Fig. 2b). Figs. 2c&d demonstrate live dye labeling of a U87 GBM xenograft slice with Cell Tracker dyes and the nuclear dye Hoechst.

### Microfluidic device fabrication through CO_2_ micromachining

Fabricating each of the components of the device required extensive CO_2_ laser calibration to achieve the desired dimensions and microfluidic architecture. The fabrication process of the platform consisted of CO_2_ laser micromachining, post-ablation processing, thermal fusion and solvent bonding, hydrophilization, and sterilization (Fig. 3). We optimized the power and speed settings of the CO_2_ laser for the fabrication of each platform component. For the microfluidic channel network, we determined different optimal CO_2_ laser power and speeds for each specific width and depth of the closed microchannels, roofless delivery channels in the central tissue culture area, and the binary network of channels that lead to a single outlet. A total of 40 microchannels, engraved onto a 300 μm-thick sheet of PMMA, lead to 40 parallel roofless delivery channels. Creation of the roofless delivery channels required multiple, shallow engravings in an iterative process to prevent polymer reflow. We found that high-power laser ablation to create microchannels with a high depth-to-width aspect ratio led to polymer reflow, resulting in channel occlusion/collapse. Additionally, high laser powers induced thermal stresses that warped the walls of the roofless delivery channels. Engraving and cutting the channel network layer took 7 min, while cutting the sealing layer took less than a minute. After successfully calibrating the laser system, we were able to generate a digital and easily reproducible protocol to fabricate the all components of the device.

The fabrication of the 40-well PMMA plate is subjected to very different manufacturing constraints because it is thicker (2.5 cm-thick) and high resolution is not needed. To cut through the full thickness of ∼2.5 cm thick PMMA, we utilized a more powerful CO_2_ laser system (wavelength of 10.6 μm and a power of 150 W). We performed multiple passes at maximum power and minimal speeds to create smooth through-cuts to create the wells of the well plate. Each pass had varying laser speeds from high-to-low to cut through the PMMA thickness gradually, from shallow to deeper cuts. The complete cutting process of the 40 well-plate took about 15 min. Alternatively, to streamline this process, we found it is more cost-effective to order the well plate from a CNC machining service (Nanchang Inte. Industrial Co., Ltd., ∼$5/unit).

### Post-processing and bonding

In order to improve the optical clarity and surface smoothness of the device, we performed a series of post-processing steps. Laser machining with CO_2_-based systems is known to introduce manufacturing defects and loss of clarity of polymer substrates during fabrication, but exposure to solvent vapors has been shown to remove these defects (Fig. 4a-d).^50^ The method developed by Ogilvie et al. utilizes chloroform vapor.^50^ Chloroform is an optimal solvent for surface treatment and modification of PMMA and direct exposure of chloroform vapor improves optical clarity, reduces surface roughness and enhances bonding to other PMMA substrates. DeVoe et al. studied ideal solvents for surface modification and solvent bonding of PMMA microfluidic substrates based on the Hildebrand solubility parameter, a measure of the cohesive molecular forces for both solvent and solute.^59^ If the cohesive forces for each material are similar, their molecules can readily co-exist, and dissolution of the solute will occur.^59,60^ PMMA has a solubility parameter of 20.1 (J/cm^3^)^1/2^, while methylene chloride and chloroform have solubility parameters of 20.2 (J/cm^3^)^1/2^ and 18.7 (J/cm^3^)^1/2^, respectively, making them ideal solvents for strong cohesive molecular forces.^60,61^ Prior to assembly, we exposed the channel network layer and the 40 well-plate to chloroform to remove rough/porous structures. We selected chloroform as the solvent for surface treatments based on its reported success by other groups.^50,62^ However, in principle, methylene chloride would also be an ideal solvent for PMMA surface treatments due to the similarity between its Hildebrand parameter and that of chloroform’s. During chloroform vapor treatments, a thin layer on the surface of PMMA presumably became dissolved by the vapor, which induced polymer reflow and allowed PMMA to spread out evenly.^50^ As shown in Figs. 4c&d, chloroform vapor treatment created a smooth surface, resulting in an optically-clear PMMA surface. This process also removed the bulges around the rims of the laser-engraved microchannels and prepared the channel network layer for bonding with the sealing layer.

To assemble the components of the device we followed previously reported thermal fusion bonding and solvent bonding techniques (Fig. 3b-e).^50,60^ These two bonding methods are widely used methods to create closed microchannels in thermoplastic substrates, due to relatively high bond strengths and overall simplicity of approach.^60^ We exposed the channel network and sealing layer to chloroform vapor followed by thermal fusion bonding. This process resulted in leak-proof and smooth microchannels with optical clarity. The width of the roofless delivery channels on the side that faces the porous membrane and the tissue sample was 126 ± 10 μm (Fig. 4e). The microchannels had a curved profile with a base width of 132 ± 2 μm and a height of 71 ± 5 μm (Fig. 4f). To complete the assembly, we bonded the sealed channel network to the 40 well-plate by solvent bonding. We briefly immersed the bottom of the 40-well plate in methylene chloride to solvate the bonding surface (Fig. 3d). For this step we utilized methylene chloride instead of chloroform; the Hildebrand solubility parameter of methylene chloride is closer to PMMA than that of chloroform,^60^ and it showed enhanced solvation of the bonding surface of the well-plate. After removal of excess solvent, we immediately bonded the well plate to the channel network with a press at room temperature (Fig. 3e). We believe that the compatibility of these bonding techniques with CO_2_ laser cutting/engraving augments the potential of the device to scale from laboratory prototyping to commercialization.

After assembly, the device required a hydrophilization treatment because the surface of PMMA is hydrophobic. The contact angle of the native surface is high (83°) and results in a resistive flow within the microchannels of the device.^63,64^ Oxygen plasma treatment has shown to reduce the contact angle (∼45°), causing the PMMA surface to be more hydrophilic and facilitating the microfluidic flow within the channels.^64^ Therefore, after assembly, we treated the assembled platform with oxygen plasma for hydrophilization followed by UV for sterilization.

### Flow Characterization

For any given batch of devices, we performed a flow assessment analysis to observe the uniformity of the flow rate across the 40 microchannels of the device. We characterized the flow distribution across multiple devices operated at 1.5 mL/hr (37.5 μL/hr/well) and observed an average flow rate of 40.6 ± 0.6 μL/hr/well with an average coefficient of variation of 12 ± 4% (ESI. Fig. S1). We attribute the discrepancy to the small imperfections during CO2 ablation and thermal bonding.

We also investigated the molecular transport through the membrane using fluorescein. We chose fluorescein because its molecular weight (M.W., 332 g/mol) is similar to the M.W. of small-molecule cancer drugs. We were specifically interested in understanding how the permeable PFTE membrane may affect the lateral spread of dye. The device operation begins when a porous PTFE membrane (with tissue on top) seals the roofless channels in the central culture area. This PTFE membrane is a 40 μm-thick fibrous meshwork with a functional pore size of 0.4 μm and porosity of 0.75. Thus, this highly porous membrane allows for lateral and vertical diffusion, and potentially allows for some flow as well. With our previous PDMS prototype,^39^ we determined that selective drug delivery can be achieved by alternating drug delivery channels with buffer channels. The delivery channels act as a source and their adjacent channels act as concentration sinks that prevent lateral spread between delivery channels. However, our PMMA device has substantial alterations in the channel dimensions and inter-channel distances with respect to our previous PDMS device; hence we needed to study the operation parameters under which we can confidently assert the absence of cross-talk between delivery channels. To establish the minimum flow rate at which the device can be safely utilized without contaminating neighboring delivery channels, we investigated lateral diffusion at seven different flow rates. We sealed the roofless channels with a PTFE membrane and ran 100 mM fluorescein across 6 of the 40 delivery channels, with the rest having PBS (Fig. 5a). With this arrangement, we analyzed the lateral diffusion one, two, and three channels away from the delivery channels.

Our experiments suggested that flow penetrates at least some portion of the porous membrane, since the observed lateral diffusion (visible as amount of fluorescence in adjacent channels) scaled with fluid velocity (Fig. 5b). This observation was consistent with some flow entering the membrane: at higher flow velocities there is less time for diffusion and at lower velocities there is more time for diffusion to occur. As a result, the lateral diffusion was higher downstream of the delivery channel compared to the middle and upstream locations. Our results indicated minimal fluorescein signal on the first channel adjacent to the delivery channel with flow rates above 0.4 mL/hr. Similarly, the signal two and three channels adjacent to the delivery channel was minimal and barely visible at flow rates above 0.4 mL/hr. However, we have no evidence that the flow lines entering from the bottom of the membrane reach the top of the membrane where flow could contact the tissue. These results are consistent with the results we obtained with our previous PDMS device,^39^ with similar velocities but higher flow rates given the larger cross-sectional area in the PMMA device.

To quantify the lateral spread from the delivery channels, we analyzed 3 regions of interest (upstream, middle, and downstream) at the delivery channels and its neighboring channels (3 on each side). Our results indicated minimal spread upstream of the channels compared to downstream (Fig. 5c,d). The baseline (dotted line) is established as the average fluorescence three channels away from the delivery channel at the highest flow rate (upstream and downstream). At 0.1 mL/hr, the slowest flow rate tested and the maximum spread, there was 13.0 ± 1.5% fluorescence one channel away (sink) relative to the average fluorescence in the delivery channel. There was 0.8 ± 0.1% relative fluorescence two channels away for flow rates above 0.4 mL/hr. At locations three channels away from the delivery channel, we reached baseline spread (± 0.1%) for flow rates above 0.4 mL/hr. With these experiments we concluded that the device should be operated at a flow rate above 0.4 mL/hr for minimized cross-talk; and that a buffer channel should be used between drug delivery channels for selective drug delivery. Thus, the microfluidic device should have no significant cross-contamination between delivery channels when utilized with alternating drug/buffer (source/sink, 20 each) channels at a flow rate of 0.4 mL/hr or higher. In principle, for other drug testing experiments that do not require buffer lanes, such as dose response studies, drugs could be placed adjacent to each other to generate drug response curves.

### Live tissue vertical diffusion assessment

Next, we used fluorescent dyes to demonstrate how our microfluidic device can achieve temporal and regioselective drug delivery. In these experiments, we characterized vertical diffusion of fluorescent compounds to understand how compounds diffuse into the tissue as a function of time. We used Hoechst, a blue-fluorescent DNA stain, and Doxorubicin (DOX), a red-fluorescent chemotherapy drug. After 7 days in culture, we transferred live, 250 μm-thick U87 GBM xenograft tumor slices to the culture area of the device. Next, we selectively delivered stripes of Hoechst and DOX to the tumor slices for varying periods of time (Fig. 6a). The fluorescent compounds were alternated as their fluorescent signals do not overlap, yielding the equivalent of a buffer lane in between for each compound. The compounds were added at different time points while the device was in continuous operation; all of the wells were initially filled with slice medium. Different tissue regions on the same device experiment were exposed for 1, 2, 4, or 8 hrs beginning that number of hours before the end of the experiment. At the corresponding time points we replaced the slice medium with the fluorescent compounds in the selected delivery wells. Fig. 6b illustrates a 14 μm tissue cross-section at the end of the experiment and the resulting delivery of Hoechst (blue) and DOX (red) at all time periods. As expected, fluorescent penetration (z-axis) increases with prolonged exposure times (Fig. 6c). Penetration depth (Fig. 6d,e) was quantified with a vertical profile. Our results indicated that there is a significant increase in vertical diffusion into the tissue over 8 hrs of exposure, for both Hoechst and DOX. When compared to Hoechst, DOX had an increased observable z-penetration, including beyond ∼200 μm in 8 hrs. The narrower profile of the Hoechst curve likely reflects the fact that Hoechst fluorescence represents only the dye bound to DNA and not free dye, while DOX fluorescence represents all dye (both bound and free).

Mass transport modeling can provide a quantitative framework for the optimization of device design, determination of assay times and interpretation of experimental data. To quantitatively describe the transport of the drugs/dyes within the tissue, we estimated their effective diffusion constants (D) using the experimentally obtained vertical fluorescent profiles of the dye/drug at different time-points (Fig. 6d,e). Assuming that the transport of the dye/drug is primarily driven by passive isotropic diffusion, we can estimate D by applying Fick’s second law of diffusion (Eqn. 1 in the Methods section). For our analysis, we discarded the fluorescence data from the 10 μm section of the tissue adjacent to the membrane, since some of the dye gets removed from the tissue surface during the washing steps post-staining. We fitted a sixth-order polynomial curve to determine the concentration of the drug/dye at the membrane surface. We then used linear interpolation of the experimental data to find the characteristic length (L) for each time-point (as defined in the Methods section). The value of D can be extracted independently from the four different time-points (1, 2, 4 and 8 hrs), and in a purely diffusive transport process, the D values should be the same for all time points. However, the estimated diffusion constant of Hoechst was 3.5 × 10^−14^ m^2^/s and that of DOX was 5.4 x 10^−14^ m^2^/s at 1 hr, whereas the values decreased monotonically until they reached values at 8 hrs of 0.7 × 10^−14^ m^2^/s and 1.3 × 10^−14^ m^2^/s, respectively (see ESI. Table 1 for all the values). This monotonic decrease indicates that a simple Fickian diffusion model might not adequately or accurately describe the transport of the drugs in the tissue. The binding or adsorption of the drug to the tissue matrix and cellular materials (DNA, proteins, lipids) can slow down the diffusive transport over time. In addition, tissue-surface evaporation can directionally drive convective transport of the drugs through the tissue. Furthermore, the tissue itself can biologically evolve over time – interaction with the drugs can alter its porosity and binding characteristics. A more realistic model would have to include binding reaction kinetics and convective flow through porous media. It should be straightforward to use finite-element modeling and analysis to simultaneously solve the convection-diffusion and binding kinetics equations, and perform a parameter sweep and iterative curve-fitting to estimate the physical constants from the experimentally obtained drug concentration data.

### Live imaging of multiplexed drug responses in GBM xenograft slices on device

*In vitro* functional tests on an individual’s cancer (e.g. live tissue from a biopsy) could help to predict that patient’s outcome, even without any molecular knowledge.^7^ Key improvements for these functional drug response assays would be real-time live tissue imaging and multiple orthogonal readouts that would reveal different temporal effects and mechanisms of action. In prior experiments with the microfluidic device, we evaluated drug responses to four different drugs with U87 xenograft tumors, we observed similar and reproducible response profiles independently of whether drug exposure was performed on device, or off device in parallel control experiments.^41^ In those experiments, the assay was performed at the end of drug exposure, off of the device with a single fluorescent cell death indicator.^41^

Here we explore the feasibility of on-device live tissue imaging and of multiple, simultaneous cell response readouts. Using an expanded repertoire of drugs, we treated U87 flank xenograft slice cultures on the microfluidic device between day 1 and day 3 in culture (Fig. 7). As shown in Fig. 7a, for each device we assayed each of the six drugs, a DMSO vehicle control, and a buffer negative control (8 total delivery conditions) twice in different regions of the slices (Fig. 7b). This pattern resulted in 16 total treatments per device, each separated by buffer lanes. After 48 hrs of exposure, we exchanged the solution in the drug delivery channels to solution containing the blue pan-nuclear dye Hoechst combined with either the green dead nuclear dye, SG, or with both CellEvent and RedDot 2 (green apoptosis indicator and far red dead nuclear dye).

After one hour we performed live imaging with viability dyes on the device to analyze cell death drug responses. As demonstrated in the previous dye experiments (Fig. 6), the short incubation time with the fluorescent viability dyes labels only the region closest to the membrane. Therefore, we measure viability where the drug concentration is closest to that of the applied drug solution (applied over two days), and we minimize any out of focus fluorescence signal. For the first set of experiments using Hoechst and SG (Fig. 7a,c,d), we used two different devices to compare repeatability and drug response. We obtained a similar response in cell death across all drugs with both devices (Fig. 7d). For the second set of experiments using Hoechst, CellEvent, and RedDot2, we observed a similar response pattern using the same drug panel but different cell death indicators, CellEvent and RedDot2, that measure apoptosis and general cell death, respectively (Fig. 7e). To further confirm our results, we removed the slices from the device and took both high-power images (20x) and confocal images (20×) at all delivery areas (Fig. 7f,k). To quantify apoptosis and general cell death at multiple locations of each of the delivery areas, we utilized an automated cellular analysis routine (Fig. 6l). As expected, we observed similar drug response patterns with this high-resolution nuclear count as with the simpler real-time, low resolution fluorescent intensity analysis. These drug response results also correlated with off-device drug responses seen previously.^41^

These set of studies demonstrated the versatility and potential of our microfluidic device. A complete assay, from drug delivery to live imaging (excluding confocal), could be performed on the platform. As shown, real-time imaging is feasible. As the cell death indicators are non-toxic to cells (except for Hoechst), in the future one could perform continuous or intermittent measurements during drug treatments to create a more sensitive assay. The short time-frame for drug exposure and analysis (less than 5 days) demonstrated how the platform could potentially serve as a tool for clinical decision-making after surgery by providing practical drug response information to guide patient therapeutic strategies. These studies also reflect the potential of the platform for early drug development stages.

### Semi-automated quantification of multi-parameter drug-responses in patient-derived colorectal cancer slice

To overcome a major roadblock to the application of our microfluidic platform to test clinical samples, we developed a semi-automated, quantitative method to analyze multi-parameter drug-response and cell-specific readouts by immunohistochemical staining in cross-sections taken of the slices after fixation. With many human tissues, autofluorescence makes non-fluorescent DAB (diaminobenzidine) immunostaining the approach of choice, but its quantitation is challenging and usually done manually.

Here, we present a semi-automated approach for quantitative image analysis of DAB staining for apoptosis and proliferation of a clinical tumor sample (Fig. 8a) that is easy to perform and uses free software (FIJI and CellProfiler). We used data from a previous experiment in which we used the device to test the effects of different drug regimens on tumor slices from a patient with a colorectal liver metastasis, then analyzed apoptosis by fluorescence and by DAB CC3+ staining, but without quantitation of the CC3+ staining.^41^ In that experiment, before drug exposure, patient-derived tumor slices (∼250 μm) were cultured off-device for 3 days, then tested for viability with an MTT assay. We selected viable slices for a comparative drug exposure experiment. We placed three slices on the device and treated each slice with four conditions for 48 hrs. Slices were treated with FOLFOX and FOLFIRI, two standard chemotherapy combinations for metastatic CRC (see Methods). These two regimens, describe the de Gramont protocol, based on infusion of fluorouracil (always combined with LV) plus either oxaliplatin (FOLFOX) or irinotecan (FOLFIRI), which has been a standard first-line chemotherapy for metastatic CRC.^65,66^ Prior to the tumor resection, the patient had received treatment with FOLFOX. Slices were also treated with staurosporine (STS, 10 μM) as a positive control, and DMSO (0.2%) as a vehicle control. Each slice was exposed to all four conditions at least once, with buffer channels in between. At the end of drug treatment, we delivered Hoechst through the drug delivery channels for 2 hrs to mark the areas of the tissue that were treated with drugs (Fig. 8b,c). Then we removed the slices from the device and treated the whole tissue with CellEvent to measure apoptosis, as well as with the red, dead nuclear dye, ethidium homodimer-1, to measure overall cell death (data not shown). In parallel we performed off-device drug treatments, but with longer drug treatment for 3 days instead of for 2 days. The complex heterogeneity of the tissue made it difficult to perform an initial chemosensitivity assessment from the tumor undersurface as we had done previously with the xenograft tumors. Most of the tumor slices had intense autofluorescent fibrotic stromal regions (Fig. 8c, red) likely resulting from the patient’s prior neoadjuvant chemotherapy and radiation. Previously, we performed an initial fluorescent analysis of this experiment in which we detected a significant increase of CellEvent apoptosis signal after STS, and a non-significant increase after FOLFIRI.^41^ This analysis required image processing to remove autofluorescent areas.

**Figure 8.**
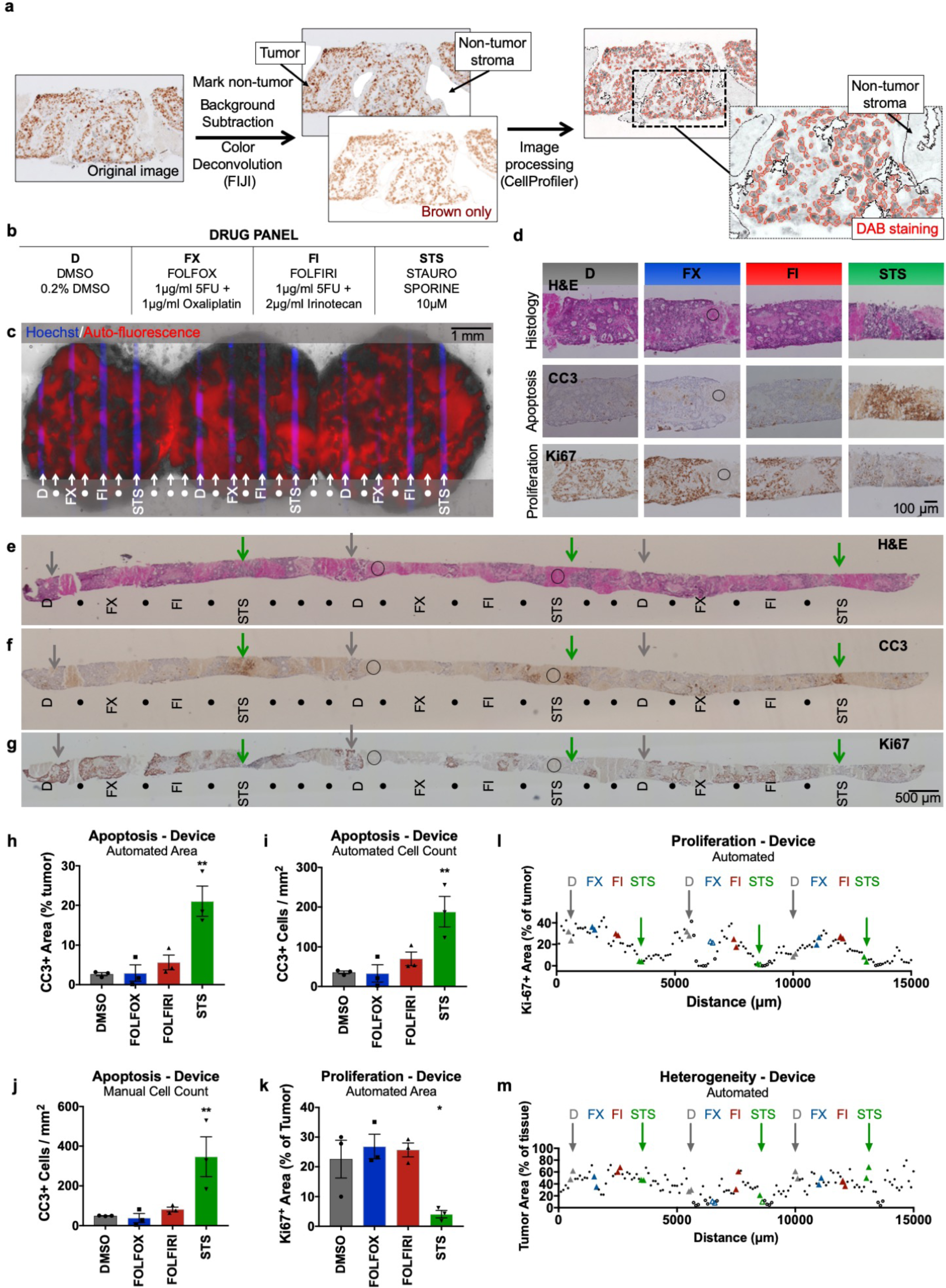
Semi-automated analysis of patient-derived liver metastasis slices from colorectal cancer (CRC) following drug exposure. Apoptosis and proliferation were quantified using immunostained metastasis slices derived from a prior experiment.^41^ (a) Workflow for image analysis of brown DAB immunohistochemical staining of tissue sections by the FIJI program followed by a custom CellProfiler routine. (b) Drug panel including vehicle control (DMSO), de Gramont protocol (FOLFOX & FOLFIRI), and staurosporine (STS). (c) CRC tumor slices after drug exposure for 2 days (off-device controls for 3 days), showing selective delivery and response to 4 different conditions (n=3 for each, labeled with Hoechst) with alternating buffer (•). Prior analysis of CellEvent live fluorescent staining of these slices demonstrated apoptosis following STS treatment^41^. (d-g) Staining of perpendicular tissue sections from the bottom halves of the CRC slices (c) shows H&E appearance and identifies apoptosis (CC3) or cell proliferation (Ki-67) for each treatment. (d) Representative 20× micrographs of drug delivery areas. (e-g) Low power micrographs of perpendicular sections show indicated locations of drug delivery channels as inferred from Hoechst staining of adjacent sections. DMSO control (D, grey arrows), STS (green arrows), and non-analyzable stromal areas (circles) are indicated for orientation. These images of CC3 and H&E staining in perpendicular sections resemble the images of staining previously seen in parallel sections taken from the top halves of the same CRC slices in Horowitz et al. Fig. 7.^41^ (h-j) Apoptosis after drug treatment based on automated area analysis (h, % of tumor), automated cell count (i, CC3+ cells/mm^2^ total area), or manual cell count (j, CC3+ cells/ mm^2^ total area) of images that largely exclude stromal (non-tumor) regions. (k,l) Proliferation after drug treatment, based on automated area analysis (% of tumor). Images were subdivided into 100-μm-wide regions for analysis along the entire tissue (l), and drug treatment locations were analyzed over a 200-μm-wide area for (k). (m) Heterogeneity of tumor area as % of total tissue, averaged from all images analyzed for each 100-μm-wide region. n=3 locations per condition, with 6 sections (>100-μm apart) analyzed per location. Ave ± SEM. One-way ANOVA versus DMSO with Dunnett’s multiple comparison test. *p<0.05, **p<0.01, ***p<0.001. Abbreviations: DAB (diaminobenzidine), HPF (high-power field).

To enable a more robust and flexible analysis for drug responses insensitive to background fluorescence, we developed quantitative assays for both apoptosis and loss of proliferation that utilize non-fluorescent immunostaining of tissue sections, the standard approach used in clinical pathology. After we sectioned the treated CRC slices, we performed CC3 immunostaining for apoptosis, Ki-67 immunostaining for proliferation, and H&E staining for histology (Figs. 8d-g). We identified the areas of drug exposure in each slice section using the Hoechst signal in adjacent sections as a guide. Then we measured drug effects by a custom image analysis routine to identify the positively stained area or cells using FIJI and CellProfiler (Fig. 8a,h-m). To validate our approach, we compared automated and manual CC3+ analyses of the same images for positive staining area and for cell counts (Fig. 8h-j). Images were taken within the 200 μm centered over the drug delivery locations, avoiding non-tumor regions. We favor the area quantitation because the automated cell counting approach sometimes undercounts adjoining cells. As seen in Figs. 8h-j, CC3 immunostaining revealed a clear apoptotic cell death response at the regions of STS treatment and no response to the other two drug combinations on-device, as well as off-device (ESI Fig. S2b). Similarly, an MTA viability analysis of slices treated off-device showed reduced viability only with STS (ESI. Fig. S2a).

To further automate DAB staining analysis, we performed analyses of proliferation and of heterogeneity across the complete tissue sections (as in Fig. 8k-m; ESI Fig. S2d,e) taken as tiled 20× images. We performed Ki67+ area quantitation on consecutive 100 μm-wide images to reveal the patterns of proliferation along all of the tissue including all drug treatment conditions (Fig. 8l). We also evaluated only the 200 μm-wide region above the drug treatment location (Fig. 8k). Ki67 immunostaining revealed reduced proliferation (as expected in areas of drug effect, the opposite of CC3) that was strong at the regions of STS treatment (Fig. 8k,l). Interestingly, off-device we observed a significant proliferation reduction for not only STS, but also for FOLFIRI, and FOLFOX, when compared to vehicle controls (ESI. Fig. S2,3). This difference likely reflects the longer treatment time off-device as compared to on-device and suggests that a longer treatment may be more sensitive for the detection drug responses. Although a longer treatment time did not result in an increase in apoptosis, it may have affected proliferation. The lack of a strong response to the chemotherapies may result from the patient’s history of previous treatment with one of the drug regimens.

This analysis also helped to quantitate and visualize the extent of tumor/stroma heterogeneity for the sampling of these CRC tumor slices. We found that our sections contained 40% ± 17% tumor (ave ± SD, range 4-80%, n=151 regions), averaged across the extent of the tissue (Fig. 8m). While the amount of tissue analyzed across the tissue remained approximately the same (ESI. Fig. S2d), the total amount of tumor analyzed per region varied, demonstrating the need for increased sampling in some regions for this tumor (ESI. Fig. S2e). These results demonstrate how this type of analysis can facilitate not only functional drug response readouts, but also address the impact of heterogeneity in clinical samples.

## Conclusions

Here we report a redesign of our microfluidic platform for functional drug testing of live tumor slices. Fabrication of the new platform is based on CO_2_ laser microfabrication and digital manufacturing of thermoplastics, techniques that are fast and inexpensive. Cost-efficient manufacturing is crucial at the pre-clinical testing stages where the platform needs to be disseminated to as many laboratories as possible. We demonstrated the functionality of the microfluidic platform with characterization studies and with multiplexed drug delivery to human GBM cell-derived xenograft slice cultures and CRC patient-derived tumor slices. We also present practical protocols for drug response readouts for cell death and proliferation, including on-device fluorescent live/dead analysis. Potential future improvements of the device could include dynamic control of flow by incorporation of microvalve multiplexers^67^ (e.g. for time-changing combinatorial mixing),^68^ micropumps,^69,70^ flow control actuators,^71^ and/or real-time mixing elements.^72^ These functional drug assays highlight the potential for future uses of our platform using clinical tumor samples for personalized medicine and utilizing intact human tissue for early stages of cancer drug development.

## Author contributions

Contributions were as follows. A.R., L.F.H., K.C., R.J.M., R.C.R., R.Y., H.K., and A.F. designed experiments. A.R., L.F.H., H.K., and K.C. performed experiments. A.R., L.F.H., K.C., N.B., G.G., and H.K. performed analysis. A.R., L.F.H., and A.F. wrote the manuscript.

## Conflicts of Interest

There are no conflicts of interest to declare.

## Acknowledgements

This work was supported by a grant from the National Cancer Institute R01 CA181445 and an International Scholars award from the Consejo Nacional de Ciencia y Tecnología de México for A.R. We would like to thank Paul Yager’s group for training and use of their laser cutting facility, Sawyer Fuller’s group for lending us their heat press equipment, and Igor Novosselov’s group for allowing us to work in their fume hood facility.

## Electronic Supplementary Information

**Supplementary Figure 1.**
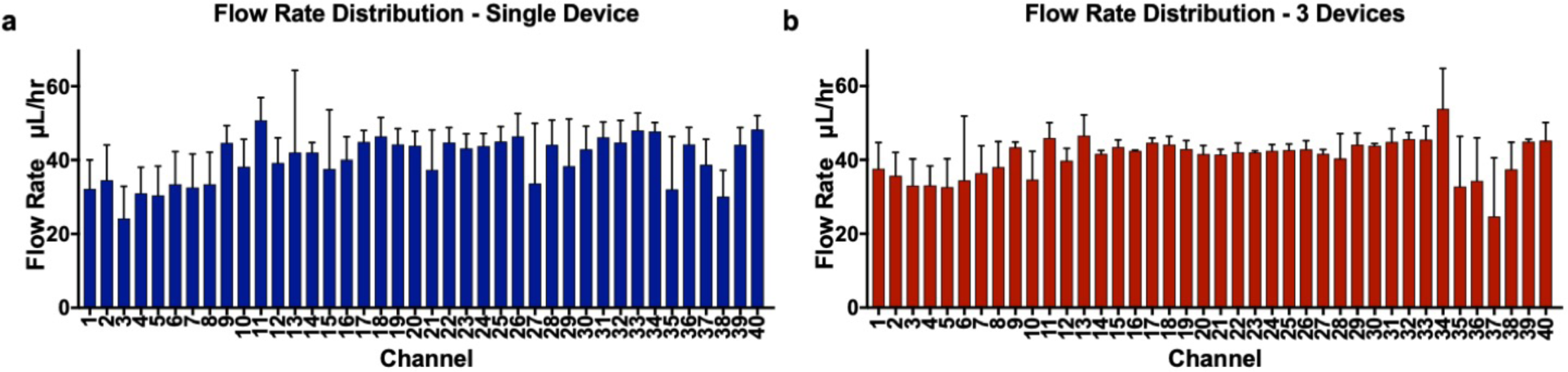
a) Flow rate distribution for a single device. A single device was operated four different times at 1.5 mL/hr (37.5 μL/hr/well) for 10 hrs (error bars show SEM, n=4). b) Flow rate distribution for a set of three devices. Each device was operated three different times at 1.5 mL/hr (37.5 μL/hr/well) for 10 hrs (error bars show SEM, n=9).

**Supplementary Figure 2.**
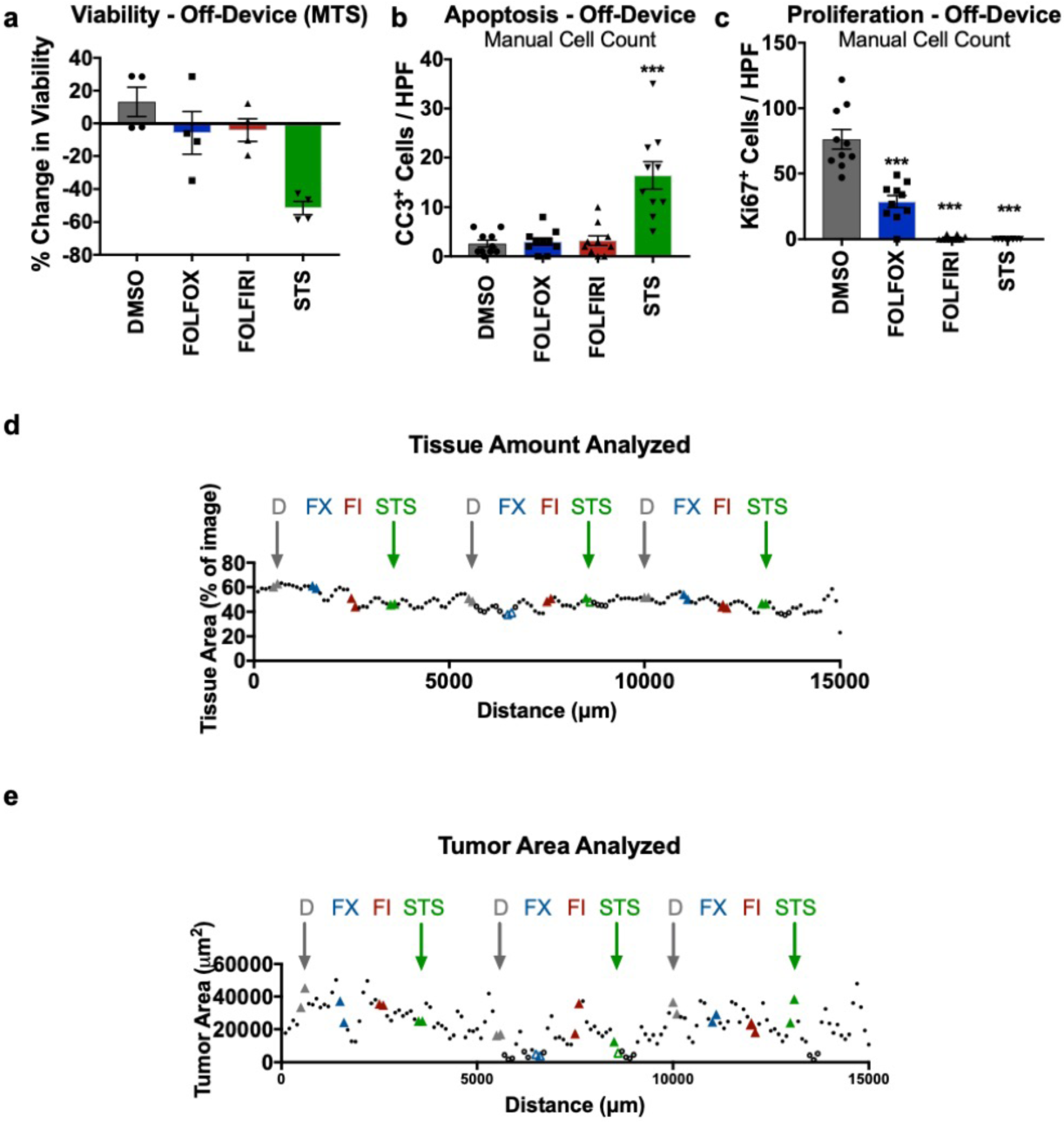
(a-c) Off-device analyses of (a) % change in viability as a function of cell metabolic activity (MTS), (b) apoptosis by manual cell count of cleaved-caspase 3 immunostaining (CC3), and (c) proliferation by manual cell count of Ki67 immunostaining for a colorectal cancer drug slices after three-day drug treatment with 1) (5FU), and Oxaliplatin, 1 μg/mL each (FOLFOX, FX), or 2) 5FU and Irinotecan, 1 μg/mL each (FOLFIRI, FI). We used DMSO as vehicle control and STS as a positive control. 4 slices per drug condition. N=4 consecutive high power fields (HPF) counted per condition. Ave ± SEM. One-way ANOVA versus DMSO with Dunnett’s multiple comparison test. *p<0.05, **p<0.01, ***p<0.001. (d,e) Tissue sampling from the automated immunostaining analysis of Ki67 in Fig. 8, with quantitation of average tissue area per image (d) and of total tumor area (e) from all 6 images used for each 100 μm-wide region. Open circles represent less than 10,000 μm^2^ for (e).

**Supplementary Figure 3.**
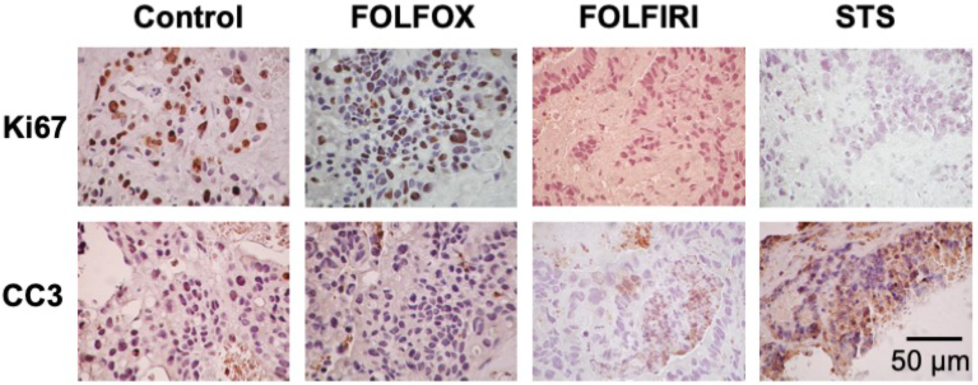
Representative 40x micrographs of perpendicular tissue sections showing proliferation (Ki67) and apoptosis (CC3) for each condition performed off-device.

**Supplementary Table 1.**
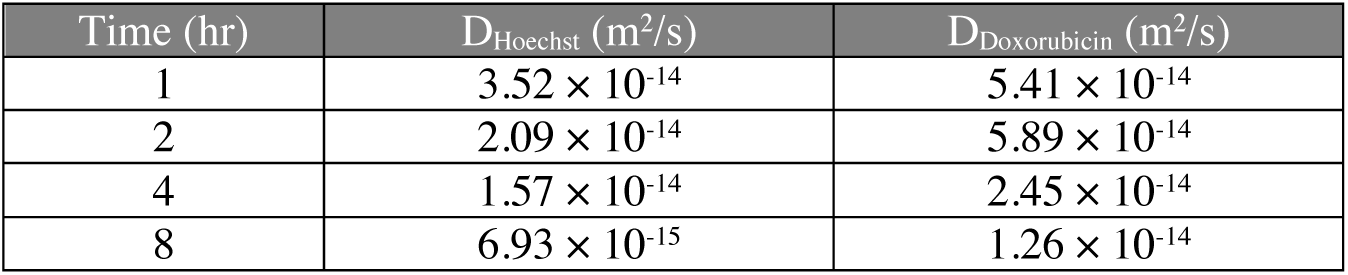
Diffusion constant values (D) for Hoechst and doxorubicin estimated (based on Eq. 1) using the experimentally obtained vertical diffusion profiles (Fig. 6) of the molecules in the U87 xenograft tumor slice at 1, 2, 4 and 8 hours.

| Time (hr) | $D_{\text{Hoechst}}$ ($\text{m}^2/\text{s}$) | $D_{\text{Doxorubicin}}$ ($\text{m}^2/\text{s}$) |
| --- | --- | --- |
| 1 | $3.52 \times 10^{-14}$ | $5.41 \times 10^{-14}$ |
| 2 | $2.09 \times 10^{-14}$ | $5.89 \times 10^{-14}$ |
| 4 | $1.57 \times 10^{-14}$ | $2.45 \times 10^{-14}$ |
| 8 | $6.93 \times 10^{-15}$ | $1.26 \times 10^{-14}$ |

